# DRD2/ANKK1 Taq1A but not COMT single nucleotide polymorphisms contribute to the link between temporal impulsivity and obesity in men

**DOI:** 10.1101/2020.06.02.130344

**Authors:** Filip Morys, Jakob Simmank, Annette Horstmann

**Affiliations:** Leipzig University Medical Centre, IFB Adiposity Diseases, 04103 Leipzig, Germany; Department of Neurology, Max Planck Institute for Human Cognitive and Brain Sciences, 04103 Leipzig, Germany; Montreal Neurological Institute, McGill University, H3A 2B2, Montreal, Canada; Department of Psychology and Logopedics, Faculty of Medicine, University of Helsinki, 00290 Helsinki, Finland; Leipzig University Medical Centre, Collaborative Research Centre 1052-A5, 04103, Leipzig, Germany

**Author notes:** **Corresponding author** Filip Morys, 3801 rue University, H3A 2B4 Montréal, Québec, Canada. **Funding** This work was funded by the Deutsche Forschungsgemeinschaft (DFG, German Research Foundation) – Projektnummer 209933838 – SFB 1052 - A05 (AH); and Federal Ministry of Education and Research (BMBF), Germany FKZ: 01E01501 (FM, JS, AH).

**Keywords:** Dopamine polymorphism, obesity, temporal impulsivity, delay discounting, DRD2, COMT, ANKK1

## Abstract

Temporal impulsivity, the tendency to choose a smaller, sooner over a larger, delayed reward, is associated with single nucleotide polymorphisms (SNPs) in COMT and DRD2-related ANKK1 genes, whose products regulate dopaminergic transmission in the brain. Temporal impulsivity is also consistently associated with obesity, sometimes in a gender-dependent fashion. Further, there seems to be no direct association between these SNPs and obesity. In this study, we investigated an interaction between BMI, COMT, and DRD2/ANKK1 SNPs, and temporal impulsivity. We tested three plausible models of associations between those variables: (1) genetic variability influencing BMI through temporal impulsivity and gender interactions, (2) genetic variability interacting with temporal impulsivity to influence BMI, (3) interaction of BMI and genetic variability influencing temporal impulsivity. We found evidence for the second model: in men, BMI was dependent on temporal impulsivity and the DRD2/ANKK1 SNP. It shows that increased temporal impulsivity combined with a disadvantageous DRD2/ANKK1 genotype might be a vulnerability factor for the development of obesity. Our study, even though cross-sectional, adds to the body of literature regarding the influence of the dopaminergic system on obesity measures. Our results point to a factor explaining discrepancies in results regarding associations of temporal impulsivity and BMI in women and men.

## 1 Introduction

A behavioural measure most consistently linked with high BMI and obesity is temporal impulsivity – a preference for smaller, but sooner, rather than larger, but delayed rewards that are not restricted to the food context^1–7^. Here, individuals discount the value of the delayed reward due to its temporal distance, hence this behaviour is called delay discounting. Such behaviour might lead to overconsumption of unhealthy, easily available and palatable, rewarding foods. The relationship of temporal impulsivity and BMI has been shown to be dependent on gender, where obese women discount rewards more than lean women, while this association is not present for men^1^. Previous research shows that delay discounting rates are related to different dopaminergic genetic variants and dopaminergic signalling within the brain. These differences were observed in adolescents with and without ADHD^8^, adults with and without alcohol abuse problem^9^, smokers^10^, gamblers^11^, and healthy adults^12–14^. Since obesity is related to changes within the dopaminergic system and in dopaminergic genetic variants^15–23^, it is important to test whether and how those alterations are related to temporal impulsivity. Elucidating how dopaminergic genetic variants and temporal impulsivity interact with obesity might thus shed light on mechanisms underlying the aetiology of obesity.

One of the dopaminergic genes previously related to delay discounting is the one coding enzyme catechol-O-methyltransferase (COMT^24^). This enzyme degrades dopamine and other catecholamines in the synaptic cleft. Its functions are predominantly effective in the prefrontal cortex, a brain structure related to executive control^25^. In other brain areas, such as the striatum, COMT’s actions are overshadowed by the dopamine transporter^26^. A single nucleotide polymorphism (SNP) of special interest in the context of delay discounting is the Val158Met polymorphism (rs4680). It results in a substitution of wild type valine into methionine in the COMT gene, rendering the enzyme less functional and thus relating to increased prefrontal dopamine levels^24^. Val homozygous participants, having relatively lower prefrontal dopamine levels, were shown to have increased preference for immediate rewards^9^, with this relationship being dependent on age^8,12^, and gender (^13^, but see:^11^). These studies suggest that, in adults, lower dopaminergic transmission in the prefrontal cortex might be related to increased delay discounting. This might specifically be due to lower dopaminergic transmission in the left dorsolateral prefrontal cortex in Val homozygotes^27^. With regard to obesity, COMT was shown to predict desirability of unhealthy foods (with Val homozygotes showing higher desirability), but not BMI^28,29^. With regard to gender differences, a study in rats showed that gender also affects the activity of COMT in the prefrontal cortex independent of the Val/Met phenotype^30^.

DRD2/ANKK1 Taq1a polymorphism substitutes the A2 allele with the A1 allele in the gene ~10kb downstream from the gene coding for striatal dopaminergic D2 receptor^31,32^. Specifically, this polymorphism is located in the Repeat And Kinase Domain Containing 1 gene (ANKK1). As a consequence of this SNP, availability of D2 receptors in the brain decreases^31–34^. This polymorphism and decreased D2 receptor density are related to addictive disorders and decreased cognitive functioning^15,35–40^. Regarding delay discounting, two studies in healthy participants showed that A1 allele carriers exhibit greater temporal impulsivity^10,41^, while another study in pathological gamblers showed no associations of temporal impulsivity and DRD2/ANKK1 Taq1A SNP^11^. The differences in results between those studies, however, might be accounted for by dopaminergic changes associated with pathological gambling. Previous investigations showed that decreased D2 receptor availability in the striatum is related to decreased glucose metabolism in the dorsolateral prefrontal cortex, orbitofrontal cortex and cingulate gyrus^16^. This suggests that DRD2/ANKK1, similarly to COMT, might affect temporal impulsivity through the prefrontal cortex. However, PET data in humans suggest that, in obese individuals, increased delay discounting rates might be related to a lower D2 receptor binding potential in the putamen^23^. The literature in the context of obesity, however, is inconsistent^40,42^ and a recent meta-analysis showed that there are most likely no effects of Taq1A polymorphism on BMI^43^.

The goal of this study is to investigate how dopaminergic genetic variants and delay discounting relate to BMI. Due to lack of consistency in reports investigating these associations, we tested all three feasible models of this three-way interaction. The first model implies that the relationship between dopaminergic genetic variants and BMI is mediated by temporal impulsivity (Figure 1a). The second one postulates that the genotype and temporal impulsivity interact to influence BMI (Figure 1b). This model is based on the (mixed) results showing that both genotype and impulsivity are associated with BMI, and we explore the possibility that they might interact to do so. The third model assumes that temporal impulsivity is influenced by the interaction of genotype and BMI (Figure 1c). This model is based on associations between genes and temporal impulsivity and BMI and impulsivity, and an exploration of an interaction between BMI and genotype. Because of previous studies that showed gender-dependent effects of obesity^1^ and dopaminergic genetic variants^13^ on delay discounting, and because activity of COMT might depend on gender^44^, we tested whether the three-way association is also gender-dependent. Since previous studies showed that decreased prefrontal metabolism and dopamine transmission are related to increased temporal impulsivity, we hypothesised that Met and A1 carriers, as compared to Met and A1 non-carriers, would show increased temporal impulsivity, but also increased BMI.

**Figure 1.**
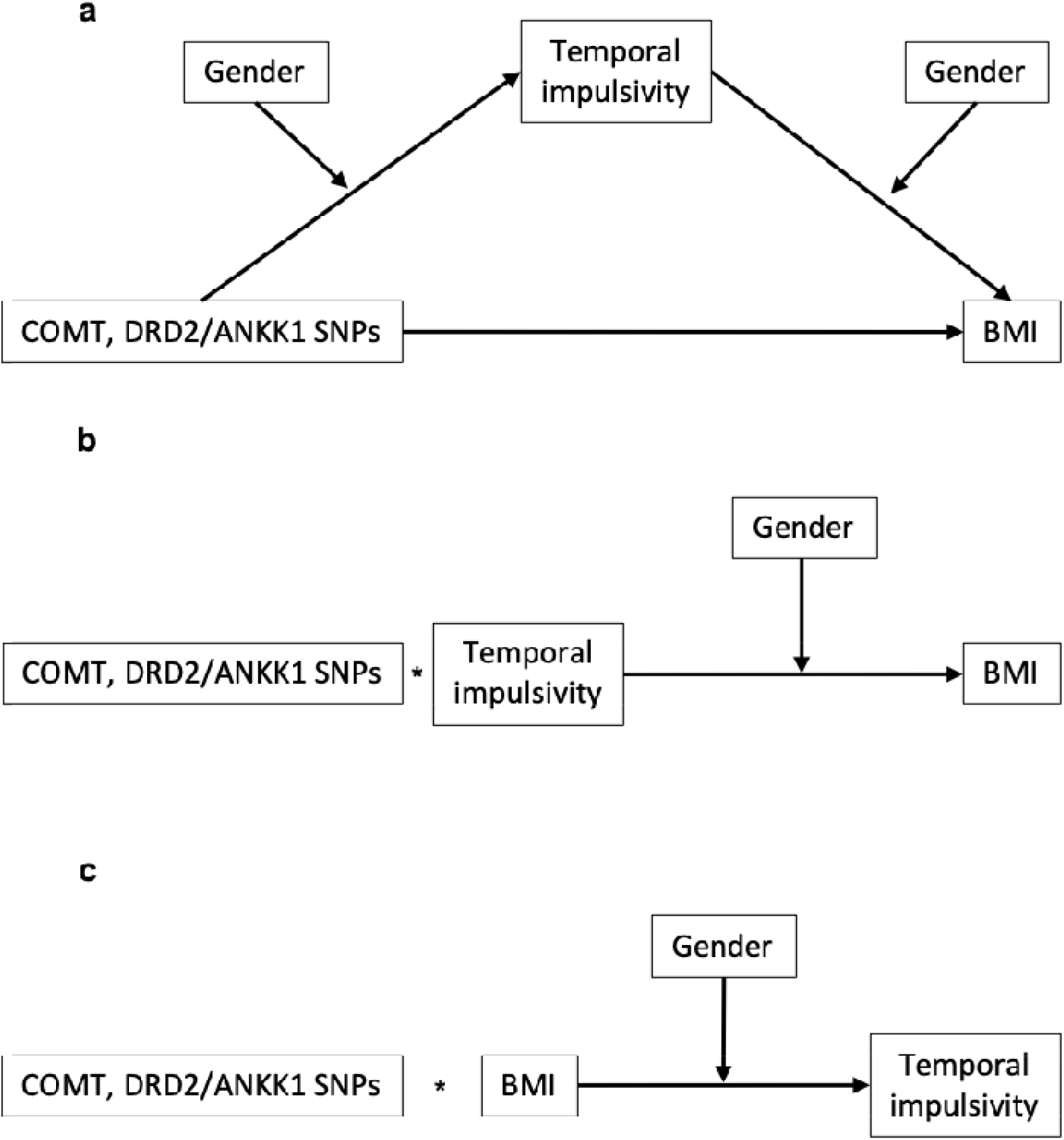
Hypothetical models of a three-way interaction between BMI, temporal impulsivity and BMI with the influence of gender that were tested in this study. **a** – dopaminergic genetic variants influence BMI through temporal impulsivity; **b** – BMI is influenced by the interaction of dopaminergic genetic variants and temporal impulsivity; **c** – temporal impulsivity is influenced by the interaction of dopaminergic genetic variants and BMI. We are unable to detect the directionality of depicted relationships, however, arrowheads were added for visualization purposes. BMI - Body Mass Index; DA – dopamine

## 2 Materials and methods

### 2.1 Participants

129 healthy lean, overweight and obese participants aged 18-35 years took part in the experiment (48 men in total; mean BMI = 27.97 kg/m^2^, SD 6.18; mean age = 26 years, SD 4 years; mean BMI of the lean group: 22.18 kg/m2, SD 1.57, overweight group: 27.37 kg/m^2^, SD 1.63, obese group: 34.20 kg/m^2^, SD 2.81). The lean group comprised 64 participants (29 men), overweight group comprised 14 participants (5 men), and the obese group comprised 51 participants (24 men). 127 Participants were at least high school graduates. Volunteers were recruited for the study with use of an Institute’s database and through posters distributed around the city of Leipzig, Germany. Participants met the following *a priori* inclusion criteria: no history of neurological/psychological diseases, no drug, cigarette or alcohol addiction, no hypertension or diabetes. Volunteers were compensated for taking part in the experiment with 7 Euro/hour. Sample in this study consists of participants recruited as parts of two different studies investigating delay discounting^45,46^, and participants recruited specifically for this study. The previous studies investigated how susceptible to environmental priming individuals with obesity are, but also focused on replicating previous associations between temporal impulsivity and BMI. Those studies implemented the same methods to test baseline delay discounting behaviours and hence can be pooled into one study. All of the studies were conducted according to the Declaration of Helsinki and approved by the Ethics Committee at the University of Leipzig. All participants gave their written informed consent prior to their participation in the studies.

### 2.2 Behavioural task

The experimental session consisted of an introduction to the experiment and a computerised delay discounting task. Detailed task procedures can be found in^45^.. During the task, participants were offered two hypothetical monetary options, a smaller one, available immediately, and a larger one, but available only after a variable delay of 1, 2, 4, 6, 9 or 12 months. We calculated and validated individual indifference points (points indicating statistical indifference between immediate and delayed options) in two separate steps. We then checked whether indifference points for ascending delays were also ascending in their monetary value (e.g. 22 Euro for 1 months, and 25 Euro for 2 months etc., instead of 22 Euro for one month, and 20 Eur for 2 months). This way we ensured that participants performed the task as instructed and expressed consistent choice behaviour. We further modelled individuals’ behaviour and calculated delay discounting parameters for each participant.

To be able to control our analyses for socioeconomic status, we have included a short questionnaire with questions regarding individual’s total income, money available to spend, satisfaction with the income, parents’ income, school degree and professional degree.

### 2.3 Genotyping

Blood samples were collected from all participants and analysed regarding their genetic information. We used Illumina Omni Express Chips (Infinium OmniExpressExome-8) to identify a number of SNPs. The analysis was done by the lab of Prof. Knut Krohn at the University of Leipzig. For this study, we extracted information regarding two specific SNPs: rs4680 – a SNP in the COMT gene (Val158Met polymorphism), and rs1800497 – a Taq1A polymorphism in the ANKK1 gene affecting expression of DRD2. Gene extraction was done using PLINK software^47^ and gwasrecode software (https://github.com/TobiWo/gwasrecode). Allele frequencies for both SNPs did not deviate from Hardy-Weinberg equilibrium (COMT: χ^2^=0.872, p=0.425; DRD2/ANKK1: χ^2^=0.483, p=0.470). Call rates for both alleles were 100%. There were no outliers regarding heterozygosity rate or genotype failure rates in the tested sample.

### 2.4 Data analysis

Behavioural data were analysed using R language within the Jupyter Notebook framework, and MATLAB 2012b (The MathWorks, Inc., Natick, Massachusetts, United States, modelling of delay discounting data using quasi-hyperbolic model). Due to relatively small sample size, we grouped participants into two groups regarding each gene – carriers and noncarriers of the Met allele, and carriers and non-carriers of the A1 allele. While the latter practice is commonly used among studies^39,41^, we did not find studies who grouped participants into Met carriers and non-carriers. However, a number of studies investigating impulsivity did not find differences between Val/Met and Met/Met individuals, enabling us to assign them into one group (^48,49^ but see:^50^). Additionally, we *post-hoc* analysed the data with three separate groups for the COMT SNP – heterozygotes and homozygotes. Similar analysis was not performed for the DRD2/ANKK1 genotype as the number of recessive homozygotes for this gene was too small to be included in such analysis.

#### 2.4.1 Delay discounting data modelling and analysis

Regarding the modelling of the delay discounting data, we followed recommendations of Simmank and colleagues^45^ who have used a sample of lean and obese participants and, to our knowledge, are the only ones to have compared the fit of hyperbolic and quasi-hyperbolic models to the delay discounting data. The hyperbolic model of delay discounting^51^ describes it as a function of a single discount factor *k*. The quasi-hyperbolic model assumes that delay discounting is dependent on two distinct parameters – beta and delta^51^. Beta describes a bias that is independent of delay – discounting the delayed reward just because it is delayed. Delta describes delay-dependent discounting – the larger the delay the more the reward is discounted. We decided to use this model since findings of Simmank and colleagues showed that the quasi-hyperbolic model fits the data better.

This model is defined by:

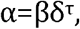

where alpha is the subjective value of a delayed reward, beta is a delay-independent bias towards immediate rewards, delta is a delay-dependent discount factor, and tau is the delay^51^. We entered choices in each individual trial into the model and beta and delta parameters were calculated (for details see ‘Estimation of Discount Function’ section in^45^). Mean adjusted R^2^ value for all participants was 81%. As an additional step, we calculated a model-independent measure of delay discounting – percentage of delayed choices. This was done to avoid exclusions of participants whose data could not be modelled.

#### 2.4.2 Data preparation

Before any analysis using delay discounting data modelling we excluded participants for whom the quasi-hyperbolic model estimation process was impossible (n=26; 10 men in total; 17 from the lean group (5 men), 2 from the overweight group (2 men), and 7 from the obese group (3 men); mean BMI = 25.63 kg/m^2^, SD 5.24; mean age = 26 years, SD 4 years), namely if the process returned beta or delta parameter values below 0 or above 1. We then tested for outliers concerning BMI, and beta and delta delay discounting parameters. *A priori* outlier exclusion criterion was as follows: values being 2.2*interquartile range below or above the first or third quartile, respectively^52–54^. One outlier regarding delta parameter was identified and excluded. This resulted in a final sample of 102 participants. Prior to all analyses, all numerical variables were z-transformed. We also checked for heteroscedasticity regarding BMI, beta and delta parameters between genetic groups. This was done using Bartlett tests. We further checked, whether genetic groups differed regarding the control variables (age and socioeconomic status), and gender distribution.

#### 2.4.3 Relationship between delay discounting and BMI

To replicate previous findings, we investigated the relationship between BMI and the DD parameters using permutation based multiple regression (lmperm package in R, 5000 permutations), while controlling for socioeconomic status. Further, we investigated the interaction of gender and BMI, because an earlier study showed gender-dependent differences in DD between obese and lean participants^1^. In this analysis, we also included age and variables describing socioeconomic status of participants. As a result, two separate regression analyses were performed, with beta and delta parameters as outcome variables, respectively (multiple comparison corrected α threshold = 0.025).

#### 2.4.4 Influence of COMT and DRD2/ANKK1 variants on delay discounting

The main goal of our study was to examine the three-way relationship between BMI, genetic variability and temporal impulsivity. One hypothesised mechanism assumed that genetic variability in COMT and DRD2/ANKK1 genes influences BMI, and that this relationship is mediated by delay discounting and moderated by gender. To test this, we first investigated whether variants of COMT and DRD2/ANKK1 genes would have an effect on delay discounting and BMI. If none of those associations proved significant, there would be no basis to conduct a formal mediation analysis. We entered genetic grouping variables and their interactions with gender into permutation-based ANCOVAs (lmperm package in R, 5000 permutations) as predictors, with age and socioeconomic status variables as covariates of no interest (control variables), and BMI, delta and beta parameters as outcome variables (3 separate ANCOVAs, multiple comparison corrected α threshold = 0.017). Here, we did not find significant relationships between genetic variability and delay discounting, or BMI (see the Results section), and decided not to perform a formal mediation analysis.

The second potential mechanism assumes that BMI is influenced by the interaction of genotype and delay discounting. To test this, we performed two permutation-based ANCOVAs with BMI as outcome measure, and genetic groups and their interactions with delta and beta parameters (hence two separate ANCOVAs) and gender as predictors. Here, age and socioeconomic status were entered as covariates of no interest (multiple comparison corrected *α* threshold = 0.025).

According to our third hypothesised mechanism temporal impulsivity is influenced by the interaction of genotype and BMI. Similarly to the second model, we performed two permutation-based ANCOVAs with delta and beta as outcome measures (hence two separate ANCOVAs), and genetic groups and their interactions with BMI and gender as predictors. Here, age and socioeconomic status were entered as covariates of no interest (multiple comparison corrected *α* threshold = 0.025).

## 3 Results

### 3.1 Genotyping

In the final sample of 102 participants, there were 4 A1 homozygotes, 71 A2 homozygotes, 32 Val homozygotes, and 24 Met homozygotes (while in the excluded sample there were 2 A1 homozygotes, 15 A2 homozygotes, 6 Val homozygotes, and 6 Met homozygotes). The minor allele frequencies in our sample were 0.17 and 0.54 for the DRD2/ANKK1 and COMT SNP, respectively (compared to 0.19 and 0.52 in a European population; based on https://www.ncbi.nlm.nih.gov/bioproject/PRJNA398795). As indicated by an ANCOVA, genetic groups (A1 carriers vs. non-carriers, and Met carriers vs. non-carriers) did not differ with regard to their age (smallest p=0.313). Chi-square tests showed that neither gender distribution nor socioeconomic status was different between groups (gender: smallest p=0.209, socioeconomic status: smallest p=0.263). We further found that variance between genetic groups regarding BMI, beta and delta parameters was homogenous, as none of Bartlett tests indicated significant differences (smallest p=0.083). Sample characteristics for investigation of percentage of delayed choices and *post-hoc* analysis dividing participants into three groups with respect to COMT genotype are presented in supplementary materials.

### 3.2 Relationship between delay discounting and BMI

We performed two regression analyses to investigate the relationship between delay discounting, BMI and gender, while controlling for age and socioeconomic status. These analyses showed that there is a significant association between BMI and delay-dependent delay discounting (regression coefficient=-0.327, p=0.017, Figure 2). Specifically, higher BMI predicted lower delta parameter (higher delay discounting behaviour). BMI did not interact with gender to influence delta parameter (regression coefficient=0.148, p=0.843). We did not find a direct association between BMI and beta parameter (regression coefficient=-0.102, p=0.347), however, the interaction of BMI and gender significantly influenced the beta parameter (regression coefficient=0.375, p=0.040, Figure 3). It shows that in women higher beta parameter (lower delay discounting) is associated with lower BMI. The opposite relationship was observed for men. This result, however, did not survive multiple comparisons correction for the number of regression analyses. Results of the analysis using percentage of delayed choices as a delay discounting measure are presented in supplementary materials. We did not find any significant associations in this investigation.

**Figure 2.**
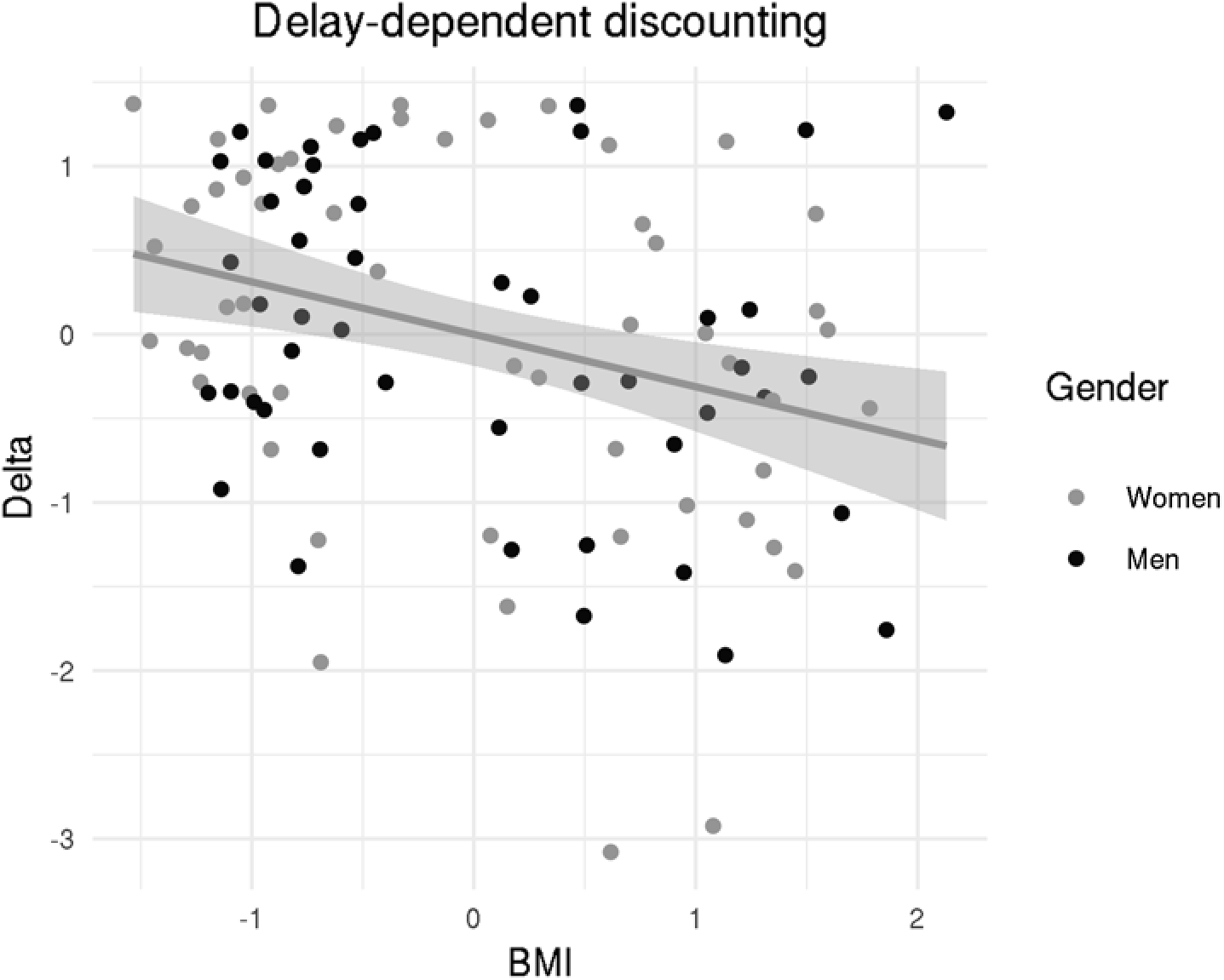
Association of BMI and delta parameter. Line indicates best fit, and shaded area represents 95% confidence intervals.

**Figure 3.**
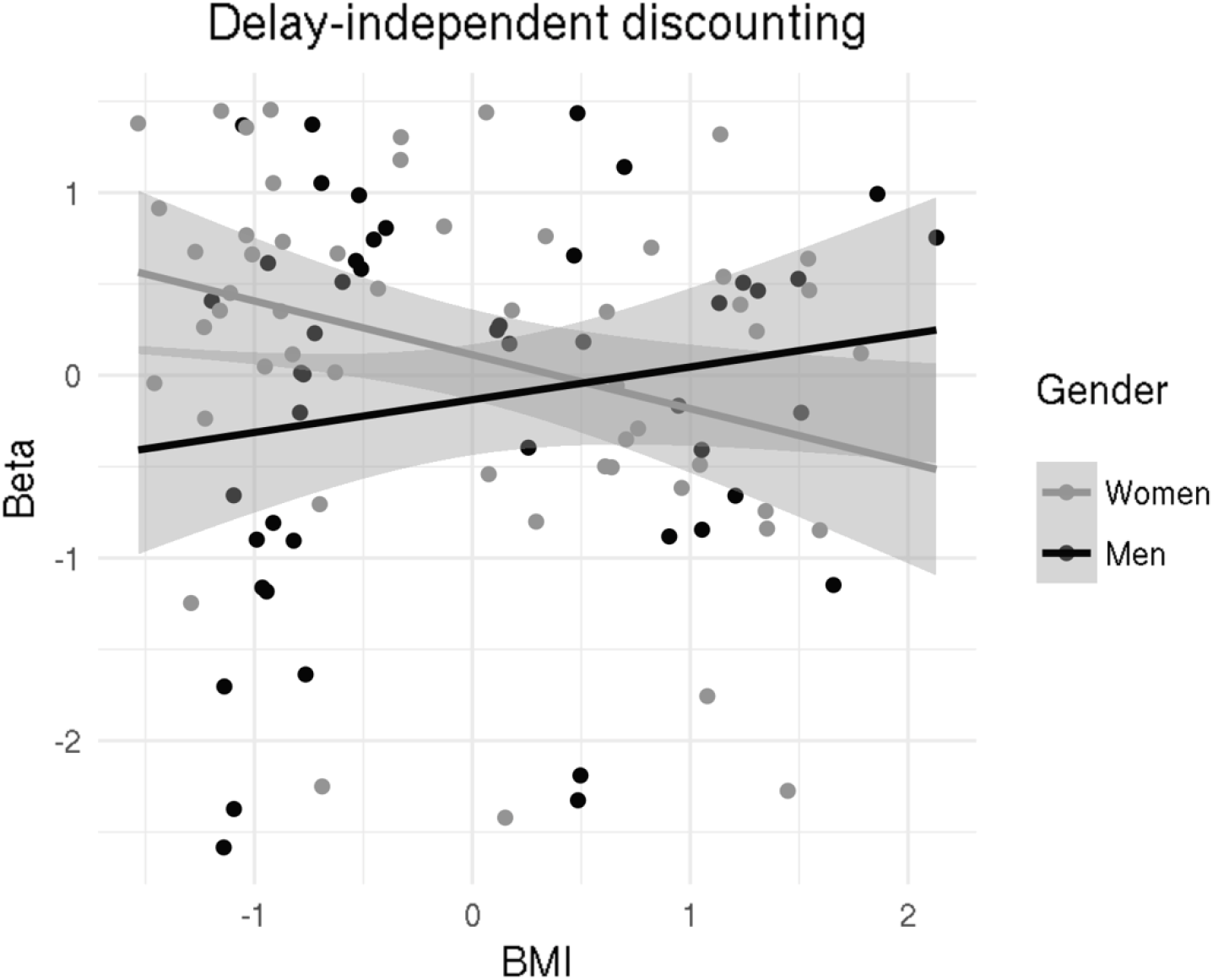
Different associations of BMI and beta parameter dependent on gender. Lines indicate best fit, and shaded areas are 95% confidence intervals. This association did not survive multiple comparisons correction (p=0.040, α=0.025).

### 3.3 Relationship between BMI, delay discounting and genetic variability in COMT and DRD2/ANKK1 genes

As a prerequisite to testing our first potential association (mediation) between the genotype, temporal impulsivity, and BMI, three separate ANCOVAs with BMI, beta and delta parameters as outcome variables were performed. None of the analyses produced significant results with regard to either DRD2/ANKK1 genetic groups, or COMT genetic groups. Interactions of genotype and gender were likewise not significant (Table 1). This indicates that we failed to find proof for a direct influence of DRD2/ANKK1 or COMT polymorphisms on delay discounting behaviour or BMI, and that gender does not play a role in this relationship. These results precluded us from conducting further mediation analysis. To test the second potential association (BMI being influenced by interaction of genotype and temporal impulsivity), we performed two separate ANCOVAs with BMI as outcome variables, and interactions of genetic groups (both groups entered in each ANCOVA), delay discounting parameters (beta and delta, resulting in two ANCOVAs), and gender as predictors. Here, we controlled for age and socioeconomic status. The analysis showed that the interaction of beta parameter, gender and DRD2/ANKK1 genotype is significantly associated with BMI (Table 2, Figure 4). Specifically, the association between BMI and beta was significantly different in men dependent on their DRD2/ANKK1 genotype. This difference was not present in women. We found no further significant associations between temporal impulsivity and COMT, or the interaction of COMT and DRD2/ANKK1 polymorphisms (Table 2).

**Table 1.** Results of ANCOVAs investigating the influence of dopaminergic genetic variants on BMI, beta, and delta delay discounting parameters. α_corrected_=0.017.

| Outcome variable | Predictors | F(1, 77) | p-value | Eta squared |
| --- | --- | --- | --- | --- |
| BMI | DRD2/ANKK1 | 0.121 | 0.667 | 0.001 |
|  | COMT | 0.003 | 1.000 | <0.001 |
|  | DRD2/ANKK1*gender | 0.013 | 0.961 | <0.001 |
|  | COMT*gender | 0.003 | 0.902 | <0.001 |
|  | DRD2/ANKK1*COMT | 0.053 | 0.843 | <0.001 |
|  | DRD2/ANKK1*COMT*gender | 0.041 | 0.921 | <0.001 |
| Beta | DRD2/ANKK1 | 0.321 | 1.000 | 0.003 |
|  | COMT | 0.631 | 0.451 | 0.006 |
|  | DRD2/ANKK1*gender | 0.540 | 0.824 | 0.005 |
|  | COMT*gender | 0.140 | 0.421 | 0.001 |
|  | DRD2/ANKK1*COMT | 0.060 | 0.706 | <0.001 |
|  | DRD2/ANKK1*COMT*gender | 1.004 | 0.442 | 0.010 |
| Delta | DRD2/ANKK1 | 3.949 | 0.166 | 0.037 |
|  | COMT | 1.899 | 0.521 | 0.018 |
|  | DRD2/ANKK1*gender | 0.115 | 1.000 | 0.001 |
|  | COMT*gender | 0.223 | 0.170 | 0.002 |
|  | DRD2/ANKK1*COMT | 3.541 | 0.062 | 0.033 |
|  | DRD2/ANKK1*COMT*gender | 0.638 | 0.706 | 0.006 |

**Table 2.** Results of ANCOVAs investigating the influence of dopaminergic genetic variants and delay discounting parameters on BMI. α_corrected_=0.025. Statistically significant values are denoted in bold.

| Outcome variable | Predictor | F(1, 69) | p-value | Eta squared |
| --- | --- | --- | --- | --- |
| BMI | DRD2/ANKK1*beta | 1.956 | 0.448 | 0.017 |
|  | DRD2/ANKK1*beta*gender | **4.065** | **0.007** | **0.034** |
|  | COMT*beta | 0.209 | 0.863 | 0.002 |
|  | COMT*beta*gender | 0.375 | 0.725 | 0.003 |
|  | DRD2/ANKK1*COMT*beta | 0.234 | 1.000 | 0.002 |
|  | DRD2/ANKK1*COMT*beta*gender | 1.193 | 0.342 | 0.010 |
| BMI | DRD2/ANKK1*delta | 2.452 | 0.882 | 0.021 |
|  | DRD2/ANKK1*delta*gender | 3.986 | 0.128 | 0.033 |
|  | COMT*delta | 0.123 | 0.281 | 0.001 |
|  | COMT*delta*gender | 0.255 | 0.583 | 0.002 |
|  | DRD2/ANKK1*COMT*delta | 0.028 | 0.765 | <0.001 |
|  | DRD2/ANKK1*COMT*delta*gender | 2.034 | 0.175 | 0.017 |

**Figure 4.**
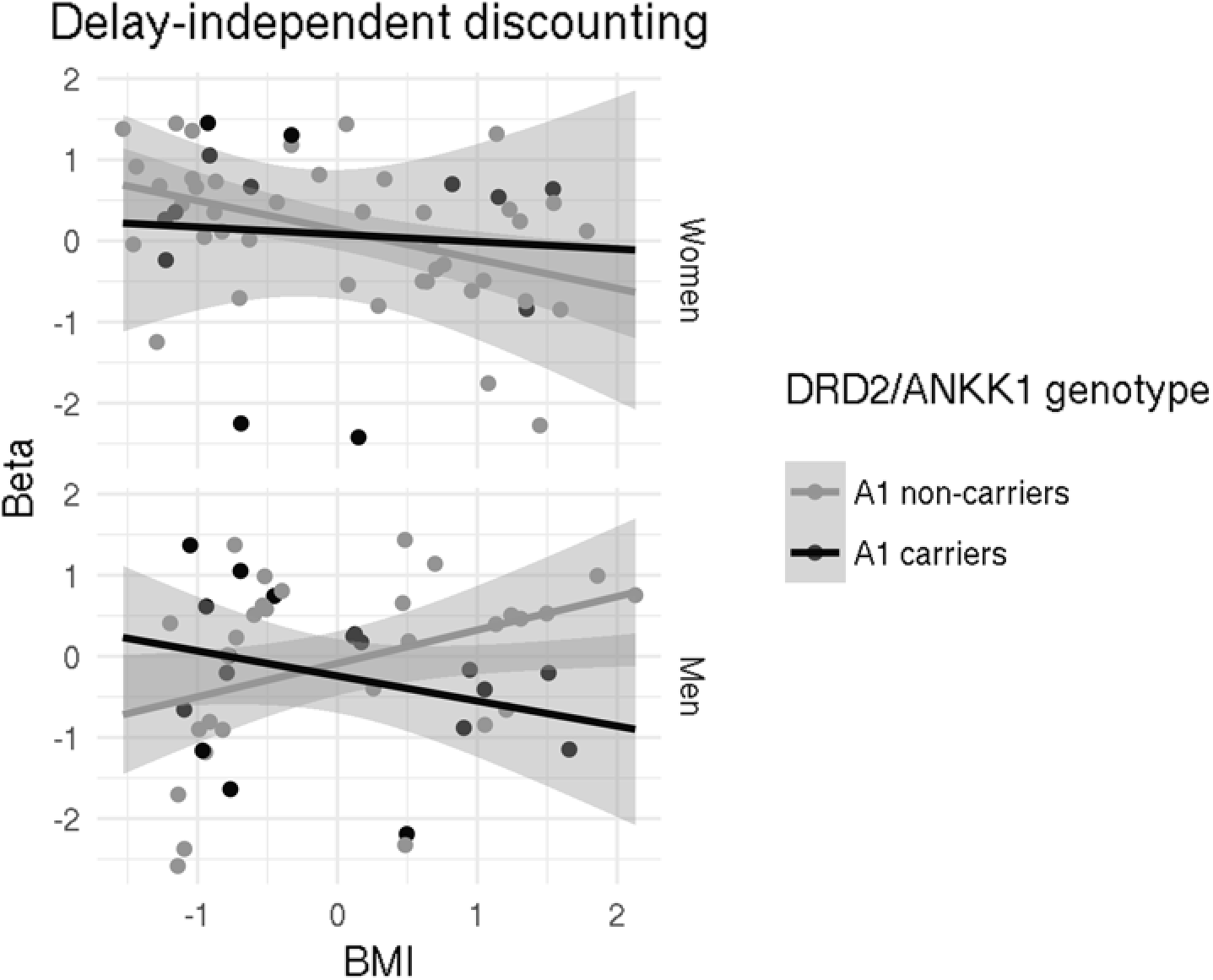
Differential associations of BMI and beta parameter dependent on the DRD2/ANKK1 Taq1A polymorphism and gender. Lines indicate best fit, and shaded areas are 95% confidence intervals.

Lastly, we tested the third potential association using two separate ANCOVAs with delta and beta parameters as outcome variables, and interactions of genetic groups (both groups entered in each ANCOVA), BMI, and gender as predictors. Here, we controlled for age and socioeconomic status. This analysis did not show any significant associations (Table 3). Results for similar analyses using percentage of delayed choices are presented in supplementary Tables S1-S3. We did not find any significant associations for this investigation. Results for *post-hoc* analysis grouping participants into three groups regarding the COMT SNP are presented in supplementary Tables S4-S6. We did not find any significant associations in this investigation.

**Table 3.** Results of ANCOVAs investigating the influence of dopaminergic genetic variants and BMI on delay discounting parameters. α_corrected_=0.025.

| Outcome variable | Predictors | F(1, 69) | p-value | Eta squared |
| --- | --- | --- | --- | --- |
| Beta | DRD2*BMI | 2.903 | 0.231 | 0.026 |
|  | DRD2/ANKK1*BMI*gender | 4.972 | 0.080 | 0.044 |
|  | COMT*BMI | 0.113 | 0.062 | <0.001 |
|  | COMT*BMI*gender | 0.003 | 0.150 | <0.001 |
|  | DRD2/ANKK1*COMT*BMI | 3.447 | 0.119 | 0.030 |
|  | DRD2/ANKK1*COMT*BMI*gender | 2.611 | 0.120 | 0.023 |
| Delta | DRD2/ANKK1*BMI | 0.227 | 0.280 | 0.002 |
|  | DRD2/ANKK1*BMI*gender | 4.189 | 0.026 | 0.035 |
|  | COMT*BMI | 1.507 | 0.638 | 0.013 |
|  | COMT*BMI*gender | 0.091 | 0.941 | <0.001 |
|  | DRD2/ANKK1*COMT*BMI | 0.019 | 0.922 | <0.001 |
|  | DRD2/ANKK1*COMT*BMI*gender | 1.038 | 0.178 | 0.009 |

## 4 Discussion

In this study we investigated the relationship between temporal impulsivity and BMI. Here, we found that higher delay-dependent delay discounting is related to increased BMI in both women and men. Further, we found that higher delay-independent delay discounting in women is related to increased BMI, with an opposite association in men. The main goal of this study, however, was to investigate the relationship between dopaminergic genetic variants (DRD2/ANKK1 and COMT polymorphisms; groups of Met carriers and noncarriers, and A1 carriers and non-carriers), temporal impulsivity and BMI. This was done to improve our understanding of the mechanisms underlying the aetiology of obesity. After probing the relationship of DRD2/ANKK1 and COMT with BMI and temporal impulsivity, we tested three different models of this relationship, where 1) temporal impulsivity mediates the relationship between dopaminergic genetic variants and BMI, 2) dopaminergic genetic variants interact with temporal impulsivity to influence BMI, 3) dopaminergic genetic variants interacts with BMI to influence temporal impulsivity. Our results provide support only for the second model. We show that the association between temporal impulsivity (delay independent parameter) and BMI is related to DRD2/ANKK1 genotype, and gender. Here, male A1 allele carriers show higher temporal impulsivity with increasing BMI, with an opposite relationship for male A1 non-carriers. In an analysis aimed at investigating a model-independent measure of delay discounting, we did not find any significant associations between BMI and temporal impulsivity and no effects of gender or dopaminergic genotype on these associations. Furthermore, we did not find any associations between BMI, temporal impulsivity and dopaminergic genotype when participants were grouped into three groups with respect to the COMT genotype.

Previous results regarding the relationships between the DRD2/ANKK1 and COMT SNPs and temporal impulsivity, but also obesity measures, are inconsistent. While COMT Val homozygotes have been often associated with increased temporal impulsivity^9,12,13^, no association with BMI has been found^28^. However, COMT polymorphism has been associated with other behaviours putting individuals at risk for obesity, such as greater desirability for unhealthy food items^29^. Evidence for the relationship between the DRD2/ANKK1 Taq1A and temporal impulsivity is rather weak and inconsistent Regarding the DRD2/ANKK1 SNP and BMI, a recent meta-analysis shows that altogether studies do not provide support for such a relationship^43^. In that report, however, authors admit that some of the studies show significant interactions of DRD2/ANKK1 polymorphism and BMI and that these might be true under some circumstances, for example in morbidly obese participants. Sun and colleagues, however, argue that the negative results might be related to the fact that analysed studies reported results in very small samples^40^. According to the authors, the DRD2/ANKK1 polymorphism explains only a small portion of variance in BMI and studies would therefore require large sample sizes to find significant associations.

Those studies point to the complexity of the relationship between behavior, more specifically temporal impulsivity, dopaminergic functioning and BMI, which might mask simple relationships between obesity and dopaminergic genetic variants, as well as between dopaminergic genetic variants and temporal impulsivity. In light of this information, the lack of significant relationships between DRD2/ANKK1 and COMT polymorphisms and temporal impulsivity or BMI in our study is not surprising. Our results are in line with most of the previous findings. The most surprising was the lack of a relationship between delay discounting and COMT, which was shown consistently in previous studies^9,12,13^. This is especially interesting given that sex impacts COMT activity as much as the SNP under investigation in our study^30^, and yet we did not see gender differences related to COMT and temporal impulsivity. It might be because the sample size in our study was relatively small. However, previous studies by Boettiger and colleagues found similar associations already in a sample of 19 participants^9^. This, however, could be explained by sample differences between the studies (alcohol dependent patients and healthy controls vs. healthy participants with a wide range of BMI). It is possible that in our case the effect size was smaller than the one in Boettiger et al. (2007), because it was masked by effects of BMI (even if those were not significant). This smaller effect size would then require a much larger sample size to detect it.

Our analyses concerning the DRD2/ANKK1 polymorphism showed that BMI is associated with an interaction of this SNP with temporal impulsivity (delay-independent parameter) in males, but not in females. Specifically, male carriers of A1 alleles who showed increased delay-independent delay discounting had increased BMI, while A1 non-carriers exhibited an opposite relationship. The delay-independent delay discounting parameter describes how much the delayed reward is discounted just because it is delayed, independent of the length of the delay. We interpret this finding as meaning that in males increased temporal impulsivity might put A1 carriers at risk of weight gain and hence lead to obesity. This finding might be related to general sex-dependent differences in DRD2 expression and function. Such differences are found in humans, where DRD2 binding potential differ between men and women^55–57^. Specifically, women show higher DRD2 binding potential in the frontal and temporal cortex, and in the thalamus^55^ and a lower binding potential in the striatum^56,57^. The lower binding potential in women in the striatum, however, is interpreted as increased dopaminergic tone^56,57^. In male rats, but not in female rats, a DRD2 antagonist eticlopride was previously shown to increase risky choice behaviours^58^. The same study showed a negative association between impulsivity and striatal DRD2 expression in male rats. Further, another study showed lower DRD2 expression in the striatum in male rats, as opposed to female rats^59^. Given that the DRD2 expression is lower in male than in female rats, that the dopaminergic tone in the striatum is lower in men than in women, and that DRD2 antagonists increase risky/impulsive choice behaviour, an even lower expression of DRD2 in men A1 carriers might exacerbate impulsive choice to a degree, where it becomes behaviourally relevant and, together with already increased temporal impulsivity, might lead to increases in BMI. A detailed interpretation of the mechanisms underlying significant findings in this study would be facilitated by applying more direct measures or manipulations of dopamine transmission in lean and obese individuals ideally investigating temporal impulsivity, such as PET experiments, dietary dopamine depletion, or using dopamine (ant-)agonists.

It is important to note that this interpretation should be viewed with caution, as our study is cross-sectional. Further investigation of the matter, ideally in a developmental dataset, is needed to confirm our interpretation. Nonetheless, this finding might help to elucidate discrepancies in results investigating relationships of BMI (or obesity) and temporal impulsivity related to gender^1,4,5,60,61^. A seminal study showing such associations is one by Weller and colleagues (2008), where increased temporal impulsivity was observed only in women with obesity, but not in men^1^. Possibly this relationship can be explained with our findings. We show that delay-independent temporal impulsivity is related to BMI in women and men in opposite directions, with this association being weaker for men. This weaker association in men is related to differential effects of the DRD2/ANKK1 polymorphism on the relationship between temporal impulsivity and BMI. Our results show that for A1 carriers and non-carriers, the relationship between temporal impulsivity and BMI is opposite. Analysing this relationship without taking genotype into account might mask the actual significant association. It is possible that this masking effect occurs in the study of Weller and colleagues^1^, and hence no association between BMI and temporal impulsivity is observed in men. Admittedly, in Weller’s study delay discounting behaviour was modelled using the hyperbolic model, which does not differentiate between delay-dependent and delay-independent delay discounting. However, with only one parameter present in the hyperbolic model, it is possible that parts of variance that we describe as delay-independent delay discounting were assigned to this one parameter. This would in turn influence the results in a way, where opposite effects of DRD2/ANKK1 SNP on BMI and temporal impulsivity association render this relationship non-significant if the genotype is not taken into account. The fact that other studies did find significant associations between BMI and temporal impulsivity in males might be due to different DRD2/ANKK1 genotype of the overall sample.

A potential limitation of this study is the fact that we grouped participants into A1 carriers and Met carriers, instead of analysing heterozygous and homozygous (dominant and recessive) participants as separate groups. This was done partly due to the fact that the sample size in our study was relatively low and hence certain groups would consist of less than 10 participants. However, the grouping strategy used in our study is not an uncommon one^39,62–64^, and a number of studies did not find impulsivity differences between Val/Met and Met/Met individuals, enabling us to assign them into one group^48,49^ (but see:^50^). Yet, to alleviate the concern that such grouping strategy might have affected our results, we performed additional analysis while grouping participants into three groups based on the COMT SNP. Here, we did not find any significant associations. This effect might occur because of insufficient sample sizes of COMT groups. In this analysis the interaction between gender, temporal impulsivity, DRD2/ANKK1 genotype and BMI was not significant. This might again be caused by insufficient sample size for such analysis with too little degrees of freedom. Further, in our primary analysis, the number of individuals in A1 and Met carriers were not equal to A1 and Met non-carriers. This was caused by the fact that we recruited participants randomly and not based on their genotype. However, in the statistical analysis we undertook steps that need to be applied when comparing two groups of unequal sizes. Additionally, due to the sample size limitations we cannot conclusively rule out the two models for which we found no significant associations between BMI, temporal impulsivity and dopaminergic genetic variants. It is possible that the effect sizes for those associations are small enough to not be detected using a sample size of 102 individuals. Further investigations on the topic are needed in groups with higher sample sizes. Additionally, we did not find any associations between model-independent measures of delay discounting (percentage of delayed choices) and BMI or dopaminergic genotype. This measure, however, is a crude measure of temporal impulsivity and does not take into account the delay of choices and magnitude of offered rewards and hence might not be sensitive enough to subtle differences in delay discounting behaviour. Lastly, overweight and obesity in our sample were classified based on BMI, which is not without its limitations – for example participants with high muscle mass might be classified as obese, while in fact they are not. However, studies point to the fact that other available measures of obesity add little further information^65^. Ideally one would use multiple measures of obesity, such as BMI, waist-to-hip ration or percentage body fat. These were, however, not collected as a part of our study.

Altogether, results of our study suggest that there is no direct influence of COMT and DRD2/ANKK1 polymorphisms on temporal impulsivity, or BMI. However, we show that the well-established association between BMI and temporal impulsivity is indeed dependent on the dopaminergic genetic variants, specifically on the DRD2/ANKK1 polymorphism in males. It might point to different mechanisms by which obesity comes about, assuming that temporal impulsivity contributes to weight-gain and, in turn, obesity. Our study potentially contributes to explaining discrepancies in results between studies showing associations between BMI and temporal impulsivity in males. We further acknowledge the need to investigate this relationship in studies with larger sample sizes and ideally a longitudinal design.

## Data availability

The dataset generated and analysed during the current study are available from the corresponding author on reasonable request.

## Acknowledgments

We would like to thank Peter Kovacs, Tobias Wohland and Lucas Scheffler for sharing their invaluable expertise and practical help on the analysis of genetic data.

## Authors’ contributions

FM, JS, and AH conceptualised the study, FM and JS collected the data, FM, JS, and AH analysed the data and wrote the manuscript.

## References

1 Weller, R. E., Cook III, E. W., Avsar, K. B. & Cox, J. E. Obese women show greater delay discounting than healthy-weight women. Appetite 51, 563–569, doi:10.1016/j.appet.2008.04.010 (2008).

2 Kishinevsky, F. I. et al. fMRI reactivity on a delay discounting task predicts weight gain in obese women. Appetite 58, 582–592, doi:10.1016/j.appet.2011.11.029 (2012).

3 McClelland, J. et al. A systematic review of temporal discounting in eating disorders and obesity: Behavioural and neuroimaging findings. Neuroscience & Biobehavioral Reviews 71, 506–528, doi:10.1016/j.neubiorev.2016.09.024 (2016).

4 Amlung, M., Petker, T., Jackson, J., Balodis, I. & MacKillop, J. Steep discounting of delayed monetary and food rewards in obesity: a meta-analysis. Psychological Medicine FirstView, 1–12, doi:10.1017/S0033291716000866 (2016).

5 Jarmolowicz, D. P. et al. Robust relation between temporal discounting rates and body mass. Appetite 78, 63–67, doi:10.1016/j.appet.2014.02.013 (2014).

6 Bickel, W. K. et al. Using crowdsourcing to compare temporal, social temporal, and probability discounting among obese and non-obese individuals. Appetite 75, 82–89, doi:10.1016/j.appet.2013.12.018 (2014).

7 Sanchez-Roige, S. et al. Genome-wide association study of delay discounting in 23,217 adult research participants of European ancestry. Nature Neuroscience 21, 16–18, doi:10.1038/s41593-017-0032-x (2018).

8 Paloyelis, Y., Asherson, P., Mehta, M. A., Faraone, S. V. & Kuntsi, J. DAT1 and COMT Effects on Delay Discounting and Trait Impulsivity in Male Adolescents with Attention Deficit/Hyperactivity Disorder and Healthy Controls. Neuropsychopharmacology 35, 2414–2426, doi:10.1038/npp.2010.124 (2010).

9 Boettiger, C. A. et al. Immediate Reward Bias in Humans: Fronto-Parietal Networks and a Role for the Catechol-O-Methyltransferase 158Val/Val Genotype. The Journal of Neuroscience 27, 14383–14391, doi:10.1523/JNEUROSCI.2551-07.2007 (2007).

10 MacKillop, J. et al. Genetic influences on delay discounting in smokers: examination of a priori candidates and exploration of dopamine-related haplotypes. Psychopharmacology 232, 3731–3739, doi:10.1007/s00213-015-4029-4 (2015).

11 Gray, J. C. & MacKillop, J. Genetic basis of delay discounting in frequent gamblers: examination of a priori candidates and exploration of a panel of dopamine-related loci. Brain and Behavior 4, 812–821, doi:10.1002/brb3.284 (2014).

12 Smith, C. T. & Boettiger, C. A. Age modulates the effect of COMT genotype on delay discounting behavior. Psychopharmacology 222, 609–617, doi:10.1007/s00213-012-2653-9 (2012).

13 Elton, A., Smith, C. T., Parrish, M. H. & Boettiger, C. A. COMT Val158Met Polymorphism Exerts Sex-Dependent Effects on fMRI Measures of Brain Function. Frontiers in Human Neuroscience 11, doi:10.3389/fnhum.2017.00578 (2017).

14 Weber, S. C. et al. Dopamine D2/3-and μ-opioid receptor antagonists reduce cue-induced responding and reward impulsivity in humans. Transl Psychiatry 6, e850, doi:10.1038/tp.2016.113 (2016).

15 Wang, G.-J. et al. Brain dopamine and obesity. The Lancet 357, 354–357, doi:10.1016/S0140-6736(00)03643-6 (2001).

16 Volkow, N. D., Wang, G.-J., Fowler, J. S. & Telang, F. Overlapping Neuronal Circuits in Addiction and Obesity: Evidence of Systems Pathology. Philosophical Transactions: Biological Sciences 363, 3191–3200 (2008).

17 Horstmann, A., Fenske, W. K. & Hankir, M. K. Argument for a non-linear relationship between severity of human obesity and dopaminergic tone. Obesity Reviews, n/a-n/a, doi:10.1111/obr.12303 (2015).

18 Horstmann, A. It wasn’t me; it was my brain – Obesity-associated characteristics of brain circuits governing decision-making. Physiology & Behavior 176, 125–133, doi:10.1016/j.physbeh.2017.04.001 (2017).

19 Mathar, D., Neumann, J., Villringer, A. & Horstmann, A. Failing to learn from negative prediction errors: Obesity is associated with alterations in a fundamental neural learning mechanism. Cortex 95, 222–237, doi:10.1016/j.cortex.2017.08.022 (2017).

20 Carnell, S., Gibson, C., Benson, L., Ochner, C. N. & Geliebter, A. Neuroimaging and obesity: current knowledge and future directions. Obesity Reviews 13, 43–56, doi:10.1111/j.1467-789X.2011.00927.x (2012).

21 Berthoud, H.-R. & Morrison, C. The brain, appetite, and obesity. Annual Review of Psychology 59, 55–92, doi:10.1146/annurev.psych.59.103006.093551 (2008).

22 Stice, E., Yokum, S., Zald, D. & Dagher, A. Dopamine-based reward circuitry responsivity, genetics, and overeating. Current Topics in Behavioral Neurosciences 6, 81–93, doi:10.1007/7854_2010_89 (2011).

23 Eisenstein, S. A. et al. Insulin, Central Dopamine D2 Receptors, and Monetary Reward Discounting in Obesity. PLoS ONE 10, e0133621, doi:10.1371/journal.pone.0133621 (2015).

24 Lachman, H. M. et al. Human catechol-0-methyltransferase pharmacogenetics: description of a functional polymorphism and its potential application to neuropsychiatric disorders. Pharmacogenetics 6, 243–250 (1996).

25 Miller, E. K. & Cohen, J. D. An Integrative Theory of Prefrontal Cortex Function. Annual Review of Neuroscience 24, 167–202, doi:10.1146/annurev.neuro.24.1.167 (2001).

26 Scatton, B., Dubois, A., Dubocovich, M. L., Zahniser, N. R. & Fage, D. Quantitative autoradiography of 3H-nomifensine binding sites in rat brain. 36, 815–822, doi:10.1016/0024-3205(85)90204-8 (1985).

27 Gianotti, L. R. R., Figner, B., Ebstein, R. P. & Knoch, D. Why Some People Discount More than Others: Baseline Activation in the Dorsal PFC Mediates the Link between COMT Genotype and Impatient Choice. Frontiers in Neuroscience 6, doi:10.3389/fnins.2012.00054 (2012).

28 Need, A. C., Ahmadi, K. R., Spector, T. D. & Goldstein, D. B. Obesity is associated with genetic variants that alter dopamine availability. Annals of Human Genetics 70, 293–303, doi:10.1111/j.1529-8817.2005.00228.x (2006).

29 Wallace, D. L. et al. Genotype status of the dopamine-related catechol-O-methyltransferase (COMT) gene corresponds with desirability of “unhealthy” foods. Appetite 92, 74–80, doi:10.1016/j.appet.2015.05.004 (2015).

30 Chen, J. et al. Functional Analysis of Genetic Variation in Catechol-O-Methyltransferase (COMT): Effects on mRNA, Protein, and Enzyme Activity in Postmortem Human Brain. The American Journal of Human Genetics 75, 807–821, doi:10.1086/425589 (2004).

31 Pohjalainen, T. et al. The A1 allele of the human D2 dopamine receptor gene predicts low D2 receptor availability in healthy volunteers. Molecular Psychiatry 3, 256–260 (1998).

32 Jönsson, E. G. et al. Polymorphisms in the dopamine D2 receptor gene and their relationships to striatal dopamine receptor density of healthy volunteers. Molecular Psychiatry 4, 290–296 (1999).

33 Ritchie, T. & Noble, E. P. Association of seven polymorphisms of the D2 dopamine receptor gene with brain receptor-binding characteristics. Neurochemical Research 28, 73–82 (2003).

34 Laakso, A. et al. The A1 allele of the human D2 dopamine receptor gene is associated with increased activity of striatal L-amino acid decarboxylase in healthy subjects. Pharmacogenet Genomics 15, 387–391 (2005).

35 Young, R. M., Lawford, B. R., Nutting, A. & Noble, E. P. Advances in molecular genetics and the prevention and treatment of substance misuse: Implications of association studies of the A1 allele of the D2 dopamine receptor gene. Addictive Behaviors 29, 1275–1294, doi:10.1016/j.addbeh.2004.06.012 (2004).

36 Munafò, M. R., Timpson, N. J., David, S. P., Ebrahim, S. & Lawlor, D. A. Association of the DRD2 gene Taq1A polymorphism and smoking behavior: A meta-analysis and new data. Nicotine & Tobacco Research 11, 64–76, doi:10.1093/ntr/ntn012 (2009).

37 Noble, E. P. D2 dopamine receptor gene in psychiatric and neurologic disorders and its phenotypes. American Journal of Medical Genetics. Part B, Neuropsychiatric Genetics: The Official Publication of the International Society of Psychiatric Genetics 116B, 103–125, doi:10.1002/ajmg.b.10005 (2003).

38 Comings, D. E. et al. The dopamine D2 receptor (DRD2) as a major gene in obesity and height. Biochemical Medicine and Metabolic Biology 50, 176–185 (1993).

39 Ariza, M. et al. Dopamine Genes (DRD2/ANKK1-TaqA1 and DRD4-7R) and Executive Function: Their Interaction with Obesity. PLOS ONE 7, e41482, doi:10.1371/journal.pone.0041482 (2012).

40 Sun, X., Luquet, S. & Small, D. M. DRD2: Bridging the Genome and Ingestive Behavior. Trends in Cognitive Sciences 21, 372–384, doi:10.1016/j.tics.2017.03.004 (2017).

41 Eisenberg, D. T. A. et al. Examining impulsivity as an endophenotype using a behavioral approach: a DRD2 TaqI A and DRD4 48-bp VNTR association study. Behavioral and Brain Functions 3, 2, doi:10.1186/1744-9081-3-2 (2007).

42 Spitz, M. R. et al. Variant alleles of the D2 dopamine receptor gene and obesity. Nutrition Research 20, 371–380, doi:10.1016/S0271-5317(00)00130-5 (2000).

43 Benton, D. & Young, H. A. A meta-analysis of the relationship between brain dopamine receptors and obesity: A matter of changes in behavior rather than food addiction. Int J Obes 40, S12–S21, doi:10.1038/ijo.2016.9 (2016).

44 Babayan, A. et al. A mind-brain-body dataset of MRI, EEG, cognition, emotion, and peripheral physiology in young and old adults. Scientific Data 6, 180308, doi:10.1038/sdata.2018.308 (2019).

45 Simmank, J., Murawski, C., Bode, S. & Horstmann, A. Incidental rewarding cues influence economic decisions in people with obesity. Frontiers in Behavioral Neuroscience, 278, doi:10.3389/fnbeh.2015.00278 (2015).

46 Morys, F., Bode, S. & Horstmann, A. Dorsolateral and medial prefrontal cortex mediate the influence of incidental priming on economic decision making in obesity. Scientific Reports 8,17595, doi:10.1038/s41598-018-35834-1 (2018).

47 Purcell, S. et al. PLINK: a tool set for whole-genome association and populationbased linkage analyses. American Journal of Human Genetics 81, 559–575, doi:10.1086/519795 (2007).

48 Su, H. et al. Predictors of heroin relapse: Personality traits, impulsivity, COMT gene Val158met polymorphism in a 5-year prospective study in Shanghai, China. American Journal of Medical Genetics Part B: Neuropsychiatric Genetics 168, 712–719, doi:10.1002/ajmg.b.32376 (2015).

49 Ziegler, D. A. et al. Motor impulsivity in Parkinson disease: Associations with COMT and DRD2 polymorphisms. Scandinavian Journal of Psychology 55, 278–286, doi:10.1111/sjop.12113 (2014).

50 Leehr, E. J. et al. A Putative Association of COMT Val(108/158)Met with Impulsivity in Binge Eating Disorder. European Eating Disorders Review 24, 169–173, doi:10.1002/erv.2421 (2016).

51 Laibson, D. Golden eggs and hyperbolic discounting. The Quarterly Journal of Economics, 443–477 (1997).

52 Tukey, J. W. Exploratory Data Analysis. 1 edition edn, (Pearson, 1977).

53 Hoaglin, D. C., Iglewicz, B. & Tukey, J. W. Performance of Some Resistant Rules for Outlier Labeling. Journal of the American Statistical Association 81, 991–999, doi:10.1080/01621459.1986.10478363 (1986).

54 Hoaglin, D. C. & Iglewicz, B. Fine-Tuning Some Resistant Rules for Outlier Labeling. Journal of the American Statistical Association 82, 1147–1149, doi:10.1080/01621459.1987.10478551 (1987).

55 Pohjalainen, T., Rinne, J. O., Någren, K., SyvÄlahti, E. & Hietala, J. Sex Differences in the Striatal Dopamine D2 Receptor Binding Characteristics in Vivo. American Journal of Psychiatry 155, 768–773, doi:10.1176/ajp.155.6.768 (1998).

56 Laakso, A. et al. Sex differences in striatal presynaptic dopamine synthesis capacity in healthy subjects. Biological Psychiatry 52, 759–763, doi:https://doi.org/10.1016/S0006-3223(02)01369-0 (2002).

57 Kaasinen, V., Någren, K., Hietala, J., Farde, L. & O., R. J. Sex Differences in Extrastriatal Dopamine D2-Like Receptors in the Human Brain. American Journal of Psychiatry 158, 308–311, doi:10.1176/appi.ajp.158.2.308 (2001).

58 Georgiou, P. et al. Dopamine and Stress System Modulation of Sex Differences in Decision Making. Neuropsychopharmacology 43, 313, doi:10.1038/npp.2017.161 (2017).

59 Orendain-Jaime, E. N., Ortega-Ibarra, J. M. & López-Pérez, S. J. Evidence of sexual dimorphism in D1 and D2 dopaminergic receptors expression in frontal cortex and striatum of young rats. 100, 62–66, doi:10.1016/j.neuint.2016.09.001 (2016).

60 Rasmussen, E. B., Lawyer, S. R. & Reilly, W. Percent body fat is related to delay and probability discounting for food in humans. Behavioural Processes 83, 23–30, doi:10.1016/j.beproc.2009.09.001 (2010).

61 Bickel, W. K., Pitcock, J. A., Yi, R. & Angtuaco, E. J. C. Congruence of BOLD Response across Intertemporal Choice Conditions: Fictive and Real Money Gains and Losses. Journal of Neuroscience 29, 8839–8846, doi:10.1523/JNEUROSCI.5319-08.2009 (2009).

62 Eisenberg, D. T. et al. Examining impulsivity as an endophenotype using a behavioral approach: a DRD2 TaqI A and DRD4 48-bp VNTR association study. Behavioral and Brain Functions 3, 2, doi:10.1186/1744-9081-3-2 (2007).

63 Esposito-Smythers, C., Spirito, A., Rizzo, C., McGeary, J. E. & Knopik, V. S. Associations of the DRD2 TaqIA polymorphism with impulsivity and substance use: Preliminary results from a clinical sample of adolescents. 93, 306–312, doi:10.1016/j.pbb.2009.03.012 (2009).

64 Soeiro-De-Souza, M., Stanford, M. S., Soares, D., Machado-Vieira, R. & Alberto Moreno, R. Association of the COMT Met158 allele with trait impulsivity in healthy young adults. doi:10.3892/mmr.2013.1336 (2013).

65 Adab, P., Pallan, M. & Whincup, P. H. Is BMI the best measure of obesity? BMJ, k1274, doi:10.1136/bmj.k1274 (2018).

