## Supplementary materials for "DRD2/ANKK1 Taq1A but not COMT single nucleotide polymorphisms contribute to the link between temporal impulsivity and obesity in men"

**Running title:** Link between BMI, impulsivity and dopaminergic SNPs

**Authors:** Filip Morys^1,2,3^, Jakob Simmank^1,2^, Annette Horstmann^1,2,4,5^

^1^ Leipzig University Medical Centre, IFB Adiposity Diseases, 04103 Leipzig, Germany

^2^ Department of Neurology, Max Planck Institute for Human Cognitive and Brain Sciences, 04103 Leipzig, Germany

^3^ Montreal Neurological Institute, McGill University, H3A 2B2, Montreal, Canada

^4^ Department of Psychology and Logopedics, Faculty of Medicine, University of Helsinki, 00290 Helsinki, Finland

^5^ Leipzig University Medical Centre, Collaborative Research Centre 1052-A5, 04103, Leipzig, Germany

**Corresponding author**

Annette Horstmann, PhD

Department of Neurology, Max Planck Institute for Human Cognitive and Brain Sciences,

Stephanstrasse 1a

04103 Leipzig, Germany

**Supplementary results**

1. Analysis of percentage of delayed choices as a model-independent measure of delay discounting.

In the final sample for this analysis we included 118 participants (after outliers exclusions). The sample consisted of 5 A1 homozygotes, 80 A2 homozygotes, 36 Val homozygotes, and 29 Met homozygotes. The minor allele frequencies in our sample were 0.18 and 0.47 for the DRD2/ANKK1 and COMT SNP, respectively (compared to 0.19 and 0.52 in a European population; based on https://www.ncbi.nlm.nih.gov/bioproject/PRJNA398795). As indicated by an ANCOVA, genetic groups (A1 carriers vs. non-carriers, and Met carriers vs. non-carriers) did not differ with regard to their age (smallest p=0.243). Chi-square tests showed that neither gender distribution nor socioeconomic status was different between groups (gender: smallest p=0.104, socioeconomic status: smallest p=0.293). We further found that variance between genetic groups regarding BMI, beta and delta parameters was homogenous, as none of Bartlett tests indicated significant differences (smallest p=0.123). Allele frequencies for both SNPs did not deviate from Hardy-Weinberg equilibrium (COMT: χ^2^=1.145, p=0.274; DRD2/ANKK1: χ^2^=0.531, p=0.531).

We performed a regression analyses to investigate the relationship between percentage of delayed choices, BMI and gender, while controlling for age and socioeconomic status. This analyses showed no significant associations between BMI and percentage of delayed choices, and lack of gender interactions (smallest p=0.180).

2. Post-hoc analysis of relationship between BMI, temporal impulsivity and dopaminergic genetic variants using three groups regarding the COMT SNP.

As indicated by an ANCOVA, genetic groups (COMT heterozygotes and recessive and dominant homozygotes) did not differ with regard to their age (p=0.390). Chi-square tests showed that individual’s total income differed between groups (p=0.031), but gender distribution or other socioeconomic status variables were not different between groups (gender: p=0.990, socioeconomic status: lowest p=263). We further found that variance between genetic groups regarding BMI, beta and delta parameters was homogenous, as none of Bartlett tests indicated significant differences (smallest p=0.304).

**Supplementary tables**

Table S1 Results of ANCOVAs investigating the influence of dopaminergic genetic variants on Percentage of delayed choices.

| Outcome variable | Predictors | F(1, 90) | p-value | Eta squared |
| --- | --- | --- | --- | --- |
| Percentage of delayed choices | DRD2/ANKK1 | 0.001 | 0.997 | <0.001 |
|  | COMT | 0.186 | 0.667 | 0.002 |
|  | DRD2/ANKK1*gender | 0.805 | 0.372 | 0.007 |
|  | COMT*gender | 0.059 | 0.826 | <0.001 |
|  | DRD2/ANKK1*COMT | 1.807 | 0.182 | 0.016 |
|  | DRD2/ANKK1*COMT*gender | 0.003 | 0.957 | <0.001 |

Table S2 Results of ANCOVAs investigating the influence of dopaminergic genetic variants and percentage of delayed choices on BMI. perc_del – percentage of delayed choices

| Outcome variable | Predictor | F(1, 86) | p-value | Eta squared |
| --- | --- | --- | --- | --- |
| BMI | DRD2/ANKK1*perc_del | 0.735 | 0.394 | 0.008 |
|  | DRD2/ANKK1*perc_del*gender | 0.026 | 0.873 | <0.001 |
|  | COMT*perc_del | 0.138 | 0.711 | 0.004 |
|  | COMT*perc_del*gender | <0.001 | 0.992 | <0.001 |
|  | DRD2/ANKK1*COMT*perc_del | 2.230 | 0.139 | 0.007 |
|  | DRD2/ANKK1*COMT*perc_del*gender | 0.001 | 0.976 | <0.001 |

Table S3 Results of ANCOVAs investigating the influence of dopaminergic genetic variants and BMI on percentage of delayed choices.

| Outcome variable | Predictors | F(1, 86) | p-value | Eta squared |
| --- | --- | --- | --- | --- |
| Beta | DRD2/ANKK1*BMI | 0.765 | 0.384 | 0.007 |
|  | DRD2/ANKK1*BMI*gender | 0.526 | 0.470 | 0.006 |
|  | COMT*BMI | 2.071 | 0.154 | 0.019 |
|  | COMT*BMI*gender | 0.140 | 0.710 | 0.001 |
|  | DRD2/ANKK1*COMT*BMI | 1.470 | 0.229 | 0.014 |
|  | DRD2/ANKK1*COMT*BMI*gender | 0.281 | 0.597 | 0.003 |

Table S4 Results of ANCOVAs investigating the influence of dopaminergic genetic variants on BMI, beta, and delta delay discounting parameters. COMT grouping: dominant and recessive homozygotes, and heterozygotes.

| Outcome variable | Predictors | F(1, 73) | p-value | Eta squared |
| --- | --- | --- | --- | --- |
| BMI | DRD2/ANKK1 | 0.098 | 0.756 | 0.001 |
|  | COMT | 0.312 | 0.733 | 0.005 |
|  | DRD2/ANKK1*gender | 0.002 | 0.961 | <0.001 |
|  | COMT*gender | 0.642 | 0.529 | 0.011 |
|  | DRD2/ANKK1*COMT | 0.739 | 0.481 | 0.013 |
|  | DRD2/ANKK1*COMT*gender | 0.530 | 0.591 | 0.009 |
| Beta | DRD2/ANKK1 | 0.562 | 0.339 | 0.003 |
|  | COMT | 3.549 | 0.034 | 0.060 |
|  | DRD2/ANKK1*gender | 1.490 | 0.226 | 0.012 |
|  | COMT*gender | 0.605 | 0.549 | 0.010 |
|  | DRD2/ANKK1*COMT | 0.261 | 0.771 | 0.004 |
|  | DRD2/ANKK1*COMT*gender | 0.990 | 0.377 | 0.017 |
| Delta | DRD2/ANKK1 | 3.681 | 0.058 | 0.034 |
|  | COMT | 1.859 | 0.163 | 0.035 |
|  | DRD2/ANKK1*gender | 0.158 | 0.692 | 0.001 |
|  | COMT*gender | 0.671 | 0.514 | 0.012 |
|  | DRD2/ANKK1*COMT | 1.839 | 0.166 | 0.034 |
|  | DRD2/ANKK1*COMT*gender | 0.709 | 0.495 | 0.013 |

Table S5 Results of ANCOVAs investigating the influence of dopaminergic genetic variants and delay discounting parameters on BMI. COMT grouping: dominant and recessive homozygotes, and heterozygotes.

| Outcome variable | Predictor | F(1, 61) | p-value | Eta squared |
| --- | --- | --- | --- | --- |
| BMI | DRD2/ANKK1*beta | 1.654 | 0.203 | 0.014 |
|  | DRD2/ANKK1*beta*gender | 3.537 | 0.065 | 0.030 |
|  | COMT*beta | 1.273 | 0.287 | 0.022 |
|  | COMT*beta*gender | 1.476 | 0.237 | 0.025 |
|  | DRD2/ANKK1*COMT*beta | 1.090 | 0.343 | 0.019 |
|  | DRD2/ANKK1*COMT*beta*gender | 0.743 | 0.480 | 0.013 |
| BMI | DRD2/ANKK1*delta | 3.053 | 0.086 | 0.025 |
|  | DRD2/ANKK1*delta*gender | 4.719 | 0.128 | 0.038 |
|  | COMT*delta | 0.073 | 0.930 | 0.001 |
|  | COMT*delta*gender | 1.463 | 0.240 | 0.024 |
|  | DRD2/ANKK1*COMT*delta | 1.170 | 0.317 | 0.019 |
|  | DRD2/ANKK1*COMT*delta*gender | 0.407 | 0.667 | 0.007 |

Table S6 Results of ANCOVAs investigating the influence of dopaminergic genetic variants and BMI on delay discounting parameters. COMT grouping: dominant and recessive homozygotes, and heterozygotes.

| Outcome variable | Predictors | F(1, 61) | p-value | Eta squared |
| --- | --- | --- | --- | --- |
| Beta | DRD2/ANKK1*BMI | 4.122 | 0.046 | 0.033 |
|  | DRD2/ANKK1*BMI*gender | 4.183 | 0.045 | 0.033 |
|  | COMT*BMI | 0.334 | 0.717 | 0.005 |
|  | COMT*BMI*gender | 0.308 | 0.736 | 0.005 |
|  | DRD2/ANKK1*COMT*BMI | 3.073 | 0.053 | 0.049 |
|  | DRD2/ANKK1*COMT*BMI*gender | 1.464 | 0.239 | 0.023 |
| Delta | DRD2/ANKK1*BMI | 0.303 | 0.584 | 0.002 |
|  | DRD2/ANKK1*BMI*gender | 3.710 | 0.059 | 0.030 |
|  | COMT*BMI | 0.608 | 0.548 | 0.010 |
|  | COMT*BMI*gender | 1.755 | 0.182 | 0.029 |
|  | DRD2/ANKK1*COMT*BMI | 0.353 | 0.704 | 0.006 |
|  | DRD2/ANKK1*COMT*BMI*gender | 0.365 | 0.696 | 0.006 |
